# How Familiarity Warps Representation in Face Space

**DOI:** 10.1101/293225

**Authors:** Vassiki Chauhan, M. Ida Gobbini

## Abstract

Recognition of familiar as compared to unfamiliar faces is robust and resistant to marked image distortion or degradation. Here we tested the flexibility of familiar face recognition with a morphing paradigm where the appearance of a personally familiar face was mixed with the appearance of a stranger. The aim was to assess how categorical boundaries for recognition of identity are affected by familiarity. We found a narrower categorical boundary for the identity of personally familiar faces when they were mixed with unfamiliar identities as compared to the control condition, in which the appearance of two unfamiliar faces were mixed. Our results suggest that familiarity warps the representational geometry of face space, amplifying perceptual distances for small changes in the appearance of familiar faces that are inconsistent with the structural features that define their identities.

## Introduction

Human beings are adept at detecting, identifying and discriminating between faces, despite the high degree of visual similarity based on first-order features. A compelling explanation for how we can discriminate different identities reliably comes from the hypothesis that faces are encoded as vectors in a high multidimensional representational space (Valentine 1991; Lee et al., 2000; Leopold el al., 2001; Jiang et al., 2007). Vectors for images of the same identity are located close to each other in this multidimensional face space. Face images for different identities that are located close to each other are harder to discriminate as compared to those that are distant from each other in face space. Several studies suggest that faces we encounter in early life (Slater et al., 2010) or on a regular basis (O’Toole et al., 1994) play a dominant role in shaping the dimensions of face space. People can discriminate the identities of faces of their own race better than faces of other races (Feingold, 1914; Rhodes et al., 1989). Webster et al.(2004) showed that if images of faces from one’s own race are morphed with images of other race faces, one tends to perceive morphs of equal mixtures as the other race. The point of subjective equality for mixed ethnicity morphs is shifted by 8% to 17% towards one’s own race.

Prior research shows that faces of personally familiar identities are better discriminated and recognized than unfamiliar faces over changes in head angle, lighting, compression or squeezing across different exemplars of the same identity (Sinha et al., 2006; Harmon & Julesz, 1973; Hole et al., 2002). Clearly, the neural face system is capable of efficiently extracting the invariant identity across multiple and varied images of the same familiar individual. At the same time, small differences in images of familiar individuals can be discriminated more efficiently for familiar than for unfamiliar faces (Haxby et al., 2000; Haxby & Gobbini, 2011; Visconti di Oleggio Castello et al. 2014).

The visual appearance of a personally familiar face is learned in detail over protracted and repeated exposure during real-life interactions. In a representational space for faces the sectors populated by familiar faces may be perceptually expanded because of the rich variety of visual experiences with those faces. Much in the same way that faces of people from one’s own race are represented more richly and with more discriminating information, faces of personally familiar others are represented more richly and with more information distinguishing different views of a familiar individual, differences that carry social significance.

Here, we wanted to test how personal familiarity affects categorical changes from one facial identity to another. We created morph continua between pairs of familiar and unfamiliar identities and measured how frequently different levels of morphed faces were assigned to the original identities. Morph continua with different levels of mixture of the attributes of two stimuli are used extensively in experimental psychology to test categorical perception. Specifically, in the field of face perception, prior research using face morphs has been focused on studying how flexible categorical perception of faces is in a variety of different conditions such as age, gender and race (Angeli et al., 2008; Webster et al., 2004; Rhodes et al., 2003). Categorical perception of a stimulus refers to the idea that for a continuous range of morphs between two stimuli, there is a perception of a categorical change around the mid-point of the morph continuum, and distinctions between morph levels within each category are less distinguishable. This has been shown to be the case for with a variety of different stimuli (Harnad, 1987), highlighting how humans tend to perceive continuous variations of stimuli as discrete categories. Categorical perception of facial identity has also been reported for both familiar and unfamiliar stimuli (Beale & Keil, 1995; Rotshtein et al., 2005; Kaufmann & Schweinberger, 2004; Kikutani et al., 2008; Ramon & van Belle, 2016).

In our experiment, we wanted to assess if a shift in categorical boundary for recognizing an identity is observed when morphing a familiar and unfamiliar identity, reflecting an expansion of perceptual distances for variations among stimuli that are perceived as the familiar individual, in support of the hypothesis that exposure to personally familiarity faces alters the representational geometry of face space.

We predicted two possible outcomes. The first hypothesis poses that repeated exposures to personally familiar faces leads to the development of perceptually expanded representational subspaces for familiar individuals. Such an expansion would shift the categorical boundary between familiar and unfamiliar identities, making an equal admixture of attributes of familiar and unfamiliar individuals perceptually resemble the unfamiliar individual. This hypothesis is supported by the work of Stevenage (1998) showing that stricter criteria for naming faces of identical twins develop as a result of training, demonstrating the importance of such criteria for discriminating between very similar faces. Alternatively, under a second hypothesis, multiple exposures to faces of familiar individuals may bias the perception of an ambiguous identity (here, a morphed image) towards being labeled more easily as a familiar individual. This hypothesis is based on the evidence that different image manipulations such as squeezing or compressing images of familiar faces do not seem to disrupt the process of recognition of those identities (Sinha et al., 2006), despite the alteration of the shape of the face and the shape of the features. Therefore, the manipulation of the identity information with the use of morphs might result in a stable, enhanced recognition with a larger categorical boundary.

## Methods

### Participants

Sixteen graduate students from the Dartmouth College community participated in this experiment (5 males, 26.8 ± 2.4). Two of the participants were left-handed. For analysis, data for one participant was discarded due an error in recording responses. Therefore, the results presented in this report are from fifteen participants (5 males, 26.7 ± 2.4). Sample size was chosen based on the sample sizes used in previous reports investigating categorical perception of facial identity (Natu et al., 2016; Kaufman & Schweinberger, 2004; Jacques & Rossion, 2006; Mckone et al, 2014). All participants provided written informed consent to participate in the experiment and were compensated with cash for their time. The Dartmouth Committee for the Protection of Human Subjects approved the experiment (Protocol 21200).

### Equipment

Participants sat 50 cm away from a computer screen in a dimly lit room. The resolution of the screen was 1440 × 900 pixels. The experiment was run on a GNU/Linux workstation, and all the presentation code was written in MATLAB, using Psychophysics toolbox extensions (Brainard, 1997; Kleiner et al, 2007).

### Stimuli

The stimuli used for this experiment were grayscale pictures of three graduate students that were personally familiar to all the participants and three unfamiliar faces that were visually matched with the familiar face identities. A morph continuum between each familiar identity and its visually matched control was created with the software FantaMorph. This procedure involved placing around 150 points per image on each of the pair of face images used to create the morphs. These points lay primarily on the internal features of the face and along the silhouette of the face, as depicted in Figure 1a. The image-processing algorithm implemented in FantaMorph used these points as landmarks to align the two images. By regulating the contribution of each identity for each image, we were able to create morphs of different strengths toward one identity or the other that contributed to the morph, resulting in a morph continuum from one original face image to the other (Figure 1b). Additionally, with identical procedures as described above, three morph continuums for 6 independent sets of identities that were all unfamiliar to the participants were created to serve as controls in the experiment. For both experimental and control morph continua, two pairs were male and one pair was female. All of the original pictures for each identity were acquired in a photo studio in the laboratory with the same lighting conditions and the same distance from the camera to minimize low-level visual dissimilarities between stimuli. We also matched the luminance of all stimuli to a target luminance value (128 in RGB) in order to control for differences in visual properties of the images themselves (Willenbockel et al., 2010).

**Figure 1:**
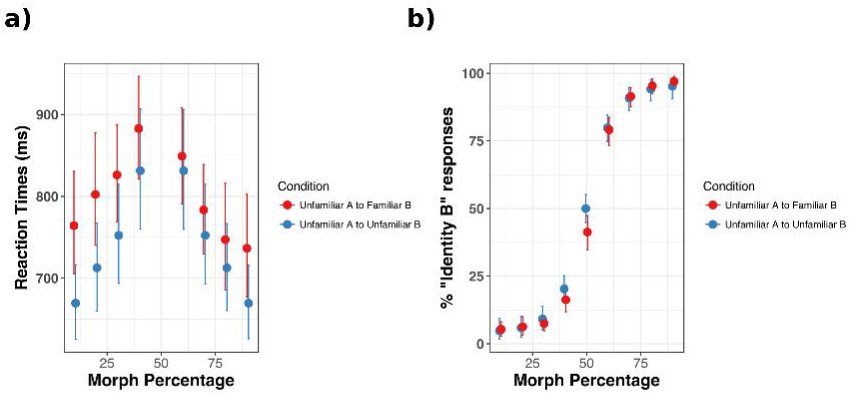
**(a)** Examples of the points chosen by FantaMorph for aligning two images. Around 150 points were chosen per face and placed around the main facial features. **(b)** Morph spectrum from an unfamiliar “Identity A” to a familiar “Identity B”. Morphs values ranged from 10% Identity B to 90% Identity B, in steps of 10%. **(c)** Example sequence from one trial of the experiment.

### Paradigm

Pictures of all the original identities used for the experiment were shown to the participants before starting the experiment. Each face was presented for 4 seconds, and participants were asked to look at the faces as they would under natural viewing conditions. Each identity was presented once.

For the experiment, each trial sequence started with the presentation of a fixation cross that remained on screen for a jittered interval between 500 and 700 ms. This was followed by the presentation of a target image for 1000 ms. This morphed image subtended 3.5 × 4.5 degrees of visual angle around the center of the screen. The target stimuli were morphed images going from 10% to 90% in steps of 10%. As soon as the target image disappeared from the screen, the two original identities from which the target morphed image was created were presented on either side of the fixation. The distance between the two original faces was 10 degrees of visual angle. The dimensions were the same for all stimuli. The two test faces stayed on screen until the subjects made a response.

The participants performed a two alternative forced choice identity recognition task. Participants were asked to respond by pressing the left or right arrow key to indicate which of the two original identities was more similar to the target face presented right before (Fig 1c). Participants were instructed to provide their response as quickly as possible, but not at the expense of accuracy.

Three blocks with the familiar/unfamiliar morphs and three blocks with unfamiliar/unfamiliar morphs were presented. Each block consisted of 108 trials, where each morphed identity was repeated 4 times at each morph percentage (10% to 90%, in steps of 10%). Thus, over the course of the entire experiment, an image from each of the identity continuum (Fig 1c) was presented to the participant 12 times. Different images from the same identity continuum were never presented in consecutive trials.

### Data Analysis

We used the R package lme4 (Bates et al., 2014) and the function ‘Anova’ from package ‘car’ for data analysis of the reaction time (RT) (Fox and Weisberg, 2011). We discarded trials that had response times shorter than 150 ms and longer than 5 s. Only correct trials were included in the analysis for RT. Trials were considered as correct when participants’ response choice matched the identity of the face that had a greater contribution to morph target. For example, if the morph was made of 40% Identity A and 60% Identity B, the correct response was considered Identity B. Therefore, this analysis excluded the 50% morph condition, since no “correct” response exists for that condition. A linear mixed model with log transformed RTs of correct trials as the dependent variable and the morph percentage of the probe and familiarity condition as independent variables was fit to the data. Scaled values of the morph percentage were used a continuous variable, whereas the familiarity condition was specified as a categorical variable with a zero sum contrast. We also included the participants, morph stimulus continuum and random intercepts for participants as random effects in the model. The RTs were log transformed in order to fit the assumptions of linear mixed models.

We also analyzed the percentage of responses for each of the face categories that the morph spectrum was built with. We refer to these responses as “Percentage Identity B” responses. In the unfamiliar-familiar morphs, “Identity B” corresponded to the familiar identity. In unfamiliar-unfamiliar morphs, the “Identity B” was an arbitrary unfamiliar face. To account for the different possible choices for labeling “Identity B” in the control condition, we adopted a bootstrapping procedure. We calculated all possible combinations for which of the two original unfamiliar identities was labeled as the “Identity B”. We then averaged responses across these different combinations and calculated 95% bootstrapped confidence intervals around the average. These are the values that have been reported in the figure and tables, and used for the statistical analyses.

For analyzing the percentage Identity B responses, a generalized linear mixed model with binomial error distribution and logit model as linking function was built. We constructed the model with the categorical response (“Identity A” or “Identity B”) as the dependent variable, morph percentage and familiarity condition as independent variables and participant, morph stimulus continuum and random intercepts for participants as random effects. Scaled values of the morph percentage were used as continuous variable, whereas the familiarity condition was specified as a categorical variable with a zero sum contrast. Statistical significance of the main effects and interaction effects was tested using a Type 3 Analysis of Deviance, as implemented in the package car.

## Data availability

Raw data and the code will be made available after acceptance at: https://github.com/vassiki/CatPercep

## Results

### Reaction Times

The estimates of the model revealed that overall, participants were slower in responding to morphs that contained identity information from familiar exemplars as compared to morphs between two unfamiliar identities (Unfamiliar-Familiar morph Mean RT = 800 ± 51 ms, Unfamiliar-Unfamiliar morph Mean RT = 742 ± 64 ms). Participants’ reaction times were also slower for the more ambiguous identities falling in the middle of the morph continuum. The main effects of familiarity and morph percentage conditions on correct, log normalized reaction times were significant (familiarity χ2(1)= 10.50, p = 0.0012) and morph percentage (χ2(1) = 33.1, p<0.001)(Figure 2; Table 1a and 1b). The interaction between the two main effects was also found to be significant (χ2(1) = 22.01, p<0.001). Estimates of the model determined by using the package lmerTest revealed that the interaction between the two main effects was driven by the presence of familiarity information for morphs along the morph continuums away from the familiar identities. Means and bootstrapped confidence intervals are included in the Tables 1a and 1b.

**Table 1a:** Average reaction times for correct trials for unfamiliar to familiar morph continuum, across different morphing percentages. Bootstrapped 95% confidence intervals are around mean reaction time.

| **Morph Percentage** | **Mean Reaction Time (ms)** | **Bootstrapped 95% CIs** |
| --- | --- | --- |
| 10% Identity B | 764 | 704, 829 |
| 20% Identity B | 802 | 740, 877 |
| 30% Identity B | 826 | 769, 885 |
| 40% Identity B | 882 | 820, 944 |
| 60% Identity B | 848 | 790, 907 |
| 70% Identity B | 783 | 728, 840 |
| 80% Identity B | 747 | 686, 816 |
| 90% Identity B | 736 | 676, 805 |

**Table 1b:**
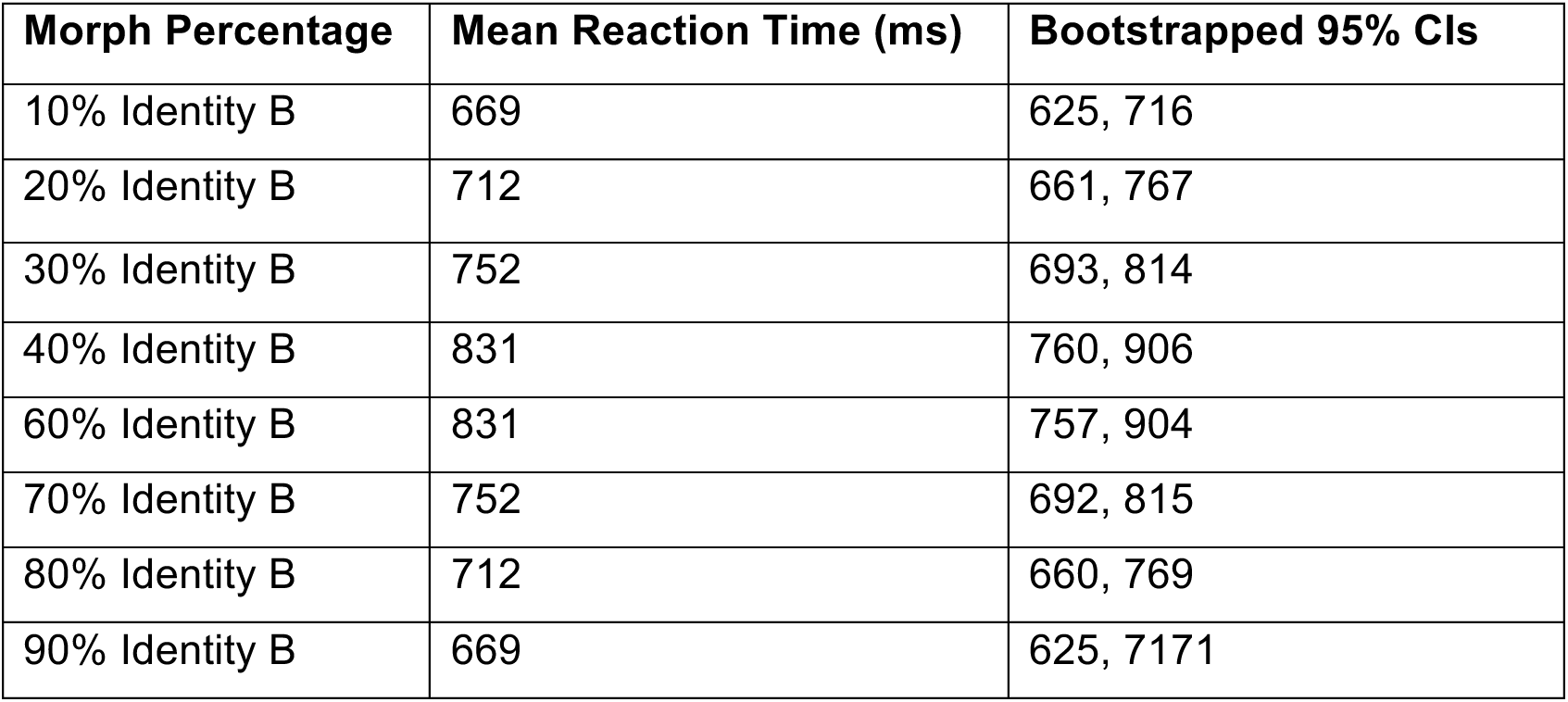
Average reaction times for correct trials for unfamiliar to unfamiliar morph continuum, across different morphing percentages. Bootstrapped 95% confidence intervals are around mean reaction time.

### Percentage Responses

Firstly, we determined the percentage of responses that corresponded to “Identity B” (Figure 1b) of the morph spectrum. “Identity B” indicates the personally familiar identity of the participants. The analysis of this dependent variable with a generalized linear mixed model revealed the main effect of morph percentage (χ2(1) = 3596.5, p<0.001) but not of familiarity condition (χ2(1) = 0.48, p=0.49). However, the interaction between morph percentage and familiarity condition was significant (χ2(1) = 4.9, p = 0.03). The mean percentage responses and bootstrapped confidence intervals are included in the Tables 2a and 2b. Unstandardized effect sizes depicting the difference in familiar and unfamiliar blocks at each morph level on percentage “Identity B” responses are included in Table 3. The effect sizes indicate that the significant interaction between the morph percentage and familiarity condition is driven by the ambiguous, 50% morph between unfamiliar and familiar identities. There is a significant reduction of 8.7%, (CI = [1.1,17.6]) in the percentage of “Identity B” responses in the unfamiliar-familiar morph condition as compared to the unfamiliar-unfamiliar morph condition.

**Table 2a:** Percentage of “Identity B” response for unfamiliar to familiar morph continuum (Identity B corresponds to the familiar identity). Bootstrapped 95% confidence intervals are around the percentage responses.

| **Morph Percentage** | **Mean Percentage Response** | **Bootstrapped 95% CIs** |
| --- | --- | --- |
| 10% Identity B | 5.4 | 2.8, 8.1 |
| 20% Identity B | 6.3 | 3.1, 10.0 |
| 30% Identity B | 7.5 | 4.8, 10.3 |
| 40% Identity B | 16.3 | 11.7, 21.5 |
| 50% Identity B | 41.3 | 34.6, 47.4 |
| 60% Identity B | 79.1 | 73.1, 83.7 |
| 70% Identity B | 91.4 | 87.6, 94.6 |
| 80% Identity B | 95.4 | 92.4, 98.0 |
| 90% Identity B | 97.0 | 94.1, 98.9 |

**Table 2b:** Percentage of “Identity B” response for unfamiliar to unfamiliar morph continuum. In these blocks, Identity B corresponded to an arbitrary unfamiliar identity. Bootstrapped 95% confidence intervals are around the percentage responses.

| **Morph Percentage** | **Mean Percentage Response** | **Bootstrapped 95% CIs** |
| --- | --- | --- |
| 10% Identity B | 4.8 | 1.7, 9.4 |
| 20% Identity B | 5.9 | 2.4, 10.2 |
| 30% Identity B | 9.2 | 5.2, 13.9 |
| 40% Identity B | 20.2 | 15.6, 25.1 |
| 50% Identity B | 50.0 | 44.6, 55.2 |
| 60% Identity B | 79.8 | 74.9, 84.4 |
| 70% Identity B | 90.8 | 86.1, 94.8 |
| 80% Identity B | 94.1 | 89.8, 97.6 |
| 90% Identity B | 95.2 | 90.6, 98.3 |

**Table 3:**
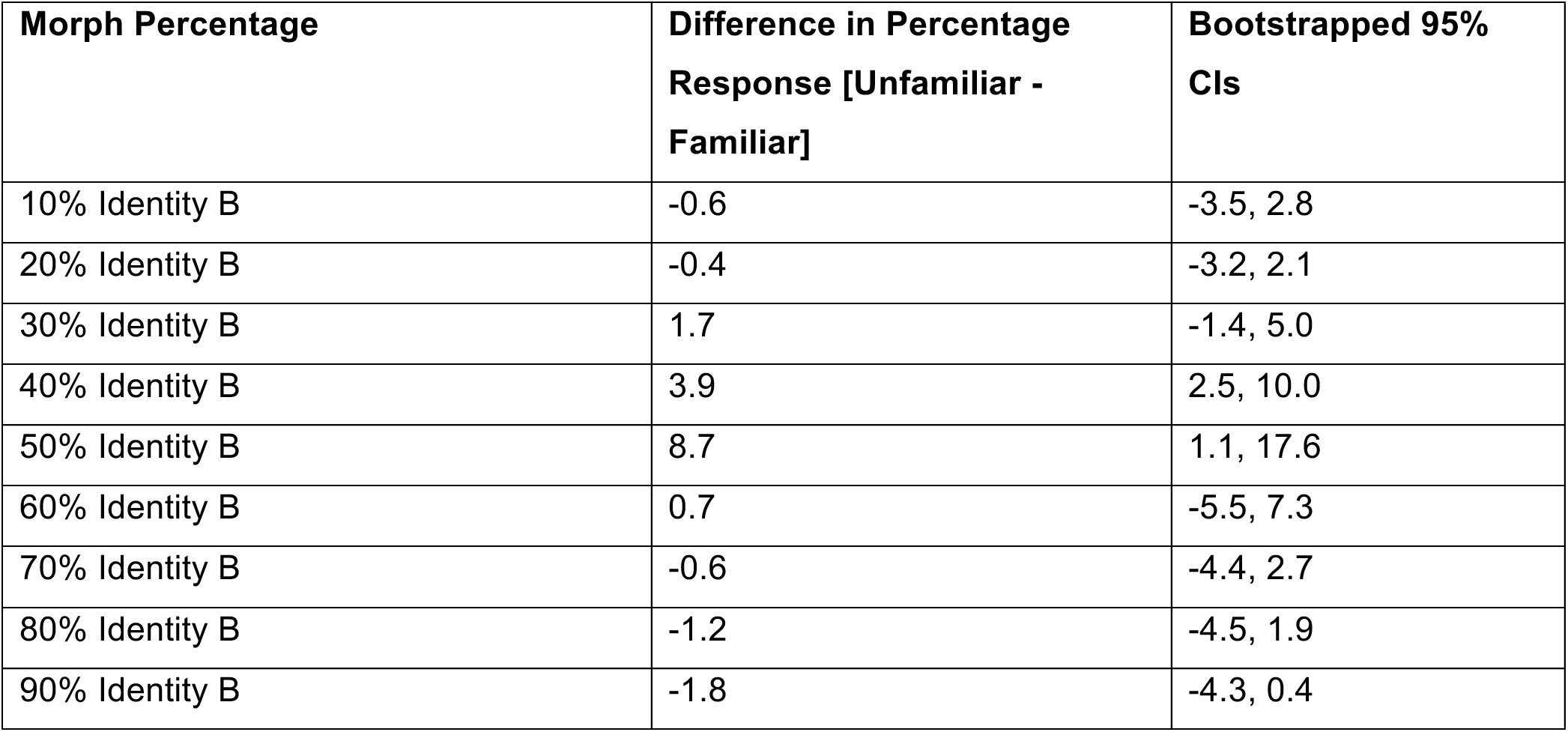
Difference in percentage “Identity B” responses in the unfamiliar-unfamiliar continuum and the unfamiliar-familiar morph continuum across different morph percentages. The third column shows bootstrapped confidence intervals around the difference in percentage responses.

| **Morph Percentage** | **Difference in Percentage Response [Unfamiliar - Familiar]** | **Bootstrapped 95% CIs** |
| --- | --- | --- |
| 10% Identity B | -0.6 | -3.5, 2.8 |
| 20% Identity B | -0.4 | -3.2, 2.1 |
| 30% Identity B | 1.7 | -1.4, 5.0 |
| 40% Identity B | 3.9 | 2.5, 10.0 |
| 50% Identity B | 8.7 | 1.1, 17.6 |
| 60% Identity B | 0.7 | -5.5, 7.3 |
| 70% Identity B | -0.6 | -4.4, 2.7 |
| 80% Identity B | -1.2 | -4.5, 1.9 |
| 90% Identity B | -1.8 | -4.3, 0.4 |

This result indicates that when the identity of morph is ambiguous with some resemblance to a familiar face, people are more conservative and less likely to respond “Identity B” when Identity B corresponds to a familiar face as compared to when both Identities A and B are unfamiliar to participants.

## Discussion

We investigated how categorical boundaries of identity are influenced by familiarity. We tested categorical decisions of identities for morph continua of face stimuli from one identity and to another. Using this experimental design, previous work has shown that perception of facial identity is ‘categorical,’ reflected in the abrupt transitions in perception of a different identity somewhere along the morph continuum (Beale & Keil, 1995; Rotshtein et al., 2005; Ramon & van Belle, 2016). Our experiment, unlike the previous work reported in the literature, tested morph continua that were created with a familiar and an unfamiliar identity. Results showed that participants were slower at categorizing morphs that were created with a familiar and an unfamiliar identity, as compared to morphs that were created by combining two unfamiliar identities. This increase in response time was greater for face morphs more like the unfamiliar individual, suggesting that the identity-inconsistent visual attributes in these morphs were more disruptive. Moreover, the categorical decision boundary was shifted toward the personally familiar face, such that the morphed image at the midpoint was more often judged to be the unfamiliar individual. This effect was demonstrated by the significant interaction between the morph level and familiarity conditions and with a significant difference between the categorical decisions for midpoints on the familiar-unfamiliar and unfamiliar-unfamiliar morph continua. While familiar faces are flexibly recognized in highly degraded or distorted images, a more conservative approach is used in labeling an ambiguous identity as a familiar individual when noise from an unfamiliar identity is added to the features of the familiar face. In the light of this result, we propose that multiple exposures to the same individuals in a variety of different viewing conditions sharpens tuning to the features that make familiar identities distinct (Tanaka et al., 1998). Conversely, an ambiguous identity is more likely to be classified as a stranger despite a strong resemblance to a familiar face since the representational geometrical space is not well-defined by learning the visual appearance of that identity through multiple exposures.

Previous research has provided evidence for categorical discrimination between identities at around the mid-point of the morph continua when two familiar identities are morphed together (Beale & Keil, 1995). When presented with two alternatives for choosing the identity of a morphed face subjects are able to choose the correct identity with high accuracy if they are familiar with the original identities (in Ramon & van Belle, 2016, identities at the extremes of the morph continua were personally familiar faces for half of the participants). In our experiment, the perception of categorical change of identity was closer to the end represented by the familiar identity indicating that when presented with an ambiguous identity, the visual system is less sensitive to changes in the features of an unfamiliar face. Our results provide strong evidence that learning through a repeated and prolonged exposure to familiar faces warps the representational geometry of face space resulting in sharper boundaries for recognition of those familiar identities. Our results are in line with research on face perception of other races. A study by Webster and colleagues (2004) showed that perception of race is influenced by the set of faces that observers are exposed to and that participants have a directional bias in determining the categorical boundary for morphs between two races (Japanese and Caucasian). The direction of this bias is determined by the participants’ race. The categorical boundary for when a face image was labeled as Japanese was closer to the Japanese face end of the morph spectrum for Japanese observers as compared to Caucasian observers, and vice versa. This result suggests that participants use a narrower boundary for categorizing exemplars from the race they belong to due higher exposure to the features of faces of that race. Similar to these results, our study shows that the categorical boundary for perceiving a face as a familiar individual is shifted towards the familiar original identity — indicating that the morphed image needs to have less noise from an unfamiliar identity to be classified as a familiar face. Learning allows a detailed representation of a face identity in different, varied visual conditions, therefore classification of identity is more conservative in terms of violation of the statistics of representation of personally familiar identities.

Our results also are in line with recent research by Faerber et al. (2016), who showed that faces of familiar international celebrities are rated as being less typical than their corresponding anti-faces, which is not true for faces of strangers (Austrian celebrities and their corresponding anti-faces). Under the face space hypothesis, the average face represents the center of the multidimensional space. The original face of the familiar celebrity is a distinct point in the multidimensional space, and its anti-face is the point that is equidistant from the average in the same space but in the opposite direction. If the subspace for familiar faces in face space is expanded, the perceptual distance between the familiar face and the population average is larger than the perceptual distance between the unfamiliar anti-face and the population average, even though these two distances are equal in terms of physical differences.

In conclusion, our experiment suggests that learning and encoding facial identity of personally familiar faces warps the multidimensional face space resulting in a representation that is more amplified, sharper, more precise and resistant to minute, ambiguity-inducing distortions that could adversely impact their recognition.

## Acknowledgements

We would like to thank Jim Haxby for his comments on a previous version of this manuscript and the Marten’s Family fund for its support.

## Author Contribution

MIG conceived the study. VC and MIG designed the experiment. VC collected and analyzed the data. VC and MIG wrote the manuscript.

---

**Figure 2: (a)** Morph percentage on the x axis and the mean reaction times (ms) on the y axis. Morph levels varied from 10% Identity B to 90% Identity B in steps of 10%. In unfamiliar-familiar blocks, “Identity B” corresponds to the familiar face. **(b)** Morph percentage on the x axis and the mean percentage “Identity B” response on the y axis. Morph levels varied from 10% Identity B to 90% Identity B in steps of 10%. In unfamiliar-familiar blocks, “Identity B” corresponds to the familiar face. In unfamiliar-unfamiliar blocks, “Identity B” is arbitrary, and the data points were calculated by bootstrapping across all possible combinations of unfamiliar-unfamiliar morph continua. The error bars represent bootstrapped 95% confidence intervals around the means.

